# Social, demographic, health care and co-morbidity predictors of tuberculosis mortality in Amazonas, Brazil: a multiple cause of death approach

**DOI:** 10.1101/658773

**Authors:** Vanderson de Souza Sampaio, Leila Cristina Ferreira da Silva, Daniel Barros de Castro, Patrícia Carvalho da Silva Balieiro, Ana Alzira Cabrinha, Antonio José Leal Costa

## Abstract

**OBJECTIVES:** To estimate TB mortality rates, describe multiple causes in death certificates in which TB was reported and identify predictors of TB reporting in death certificates in the State of Amazonas, Brazil, based on a multiple cause of death approach.

**METHODS:** Death records of residents in AM within 2006-2014 were classified based on tuberculosis reporting in the death certificate as tuberculosis not reported (TBNoR), reported as the underlying cause of death (TBUC) and as an associate cause of death (TBAC). Age standardized annual mortality rates for TBUC, TBAC and with TB reported (TBUC plus TBAC) were estimated for the State of Amazonas, using the direct standardization method and WHO 2000-2025 standard population. Mortality odds ratios (OR) of reporting TBUC and TBAC were estimated using multinomial logistic regression.

**RESULTS:** Age standardized annual TBUC and TBAC mortality rates ranged, between 5.9-7.8/10^5^ and 2.7-4.0/10^5^, respectively. TBUC was associated with residence in the State capital (OR=0.66), female sex (OR=0.87), education level (OR=0.67 and 0.50 for 8 to 11 and 12 or more school years), non-white race/skin colour (OR=1.38) and occurrence of death in the State capital (OR=1.69). TBAC was related to time (OR=1.21 and 1.22 for years 2009-11 and 2012-14), age (OR=36.1 and 16.5 for ages 15-39 and 40-64 years) and when death occurred in the State capital (OR=5.8).

**CONCLUSION:** TBUC was predominantly associated with indicators of unfavorable socioeconomic conditions and health care access constraints, whereas TBAC was mainly related to ages typical of high HIV disease incidence.

**Conflicts of interest:** None.

**Funding:** Fundação de Amparo à Pesquisa do Estado do Amazonas - FAPEAM

## Introduction

Over the last three decades tuberculosis control efforts have proven successful worldwide and by 2015, global tuberculosis incidence and mortality had fallen 18% and 47% relative to 1990 estimates [1]. Yet, tuberculosis ranks globally as the ninth leading cause of death and the leading cause from a single infectious agent [2]. In Brazil, one of the tuberculosis high burden countries according to the World Health Organization (WHO), all 2015 targets - falling incidence rate and 50% reduction in prevalence and mortality rates compared with 1990 - set in the context of the Millennium Development Goals were met [1]. Nevertheless, tuberculosis remains a public health threat nationwide with markedly heterogeneous patterns of morbidity and mortality.

The State of Amazonas has historically been among tuberculosis incidence and mortality top rank levels in Brazil. For all forms of tuberculosis, its incidence rate (70.1/100,000) ranked first in 2015 and its mortality rate (3.3/100,000) third in 2014 among Brazilian States, exceeding in 2.3 and 1.5 times the corresponding national incidence (30.9/1000,000) and mortality (2.2/100,000) estimates [3].

So far, mortality statistics based on underlying cause of death criteria have underpinned public health epidemiology analysis. A more comprehensive understanding of population mortality patterns requires knowledge of the myriad of causes related to the occurrence of death, as provided by multiple cause of death analysis [4]. In multiple cause of death analysis all causes registered in death certificates are considered, allowing enumeration of the underlying and its associated causes, including consequential causes - those following the underlying cause and directly related to death – and contributing causes – those not directly related but that, in some way, contributed to death [5].

Multiple cause of death analysis of tuberculosis provides information on what complications follow tuberculosis when assigned as the underlying cause, as well as the underlying causes when tuberculosis is reported as associate cause. It also allows a more comprehensive estimation of mortality involving tuberculosis [5].

Despite its high morbidity and mortality levels, no studies have focused on factors related to tuberculosis mortality using multiple cause of death analysis in the State of Amazonas, Brazil. Based on a multiple cause of death approach, the aims of the present study were to estimate TB mortality rates, describe multiple causes assigned in death certificates in which TB was reported and identify predictors of TB reporting in death certificates in the State of Amazonas, Brazil.

## Material and methods

Located in the north region of Brazil, Amazonas is Brazil’s largest State, with 1,559,161 km^2^, most of it covered by the Amazon forest and hydrographic basin. In 2010 it had a population of 3,483,985 inhabitants spread among 62 municipalities, 79% living in urban settings and 21% in rural areas. The average demographic density in 2010 was 2.23 inhabitants/km^2^. Only eight municipalities had more than 50,000 inhabitants - including Manaus, the State capital with 52% (1,802,014) of the State population - and in 29, the population was below 20,000. Almost 5% of the population is indigenous, mainly of Tikuna ethnicity [6].

We conducted a cross-sectional exploratory study based on a series of official mortality registers. Data was obtained from the Mortality Information System (Sistema de Informações sobre Mortalidade – SIM, available at http://datasus.saude.gov.br/) coordinated by the State of Amazonas Foundation for Public Health Surveillance (Fundação de Vigilância em Saúde do Estado do Amazonas – FVS AM). Data imputed into SIM comes from a nationwide standard death certificate filled by physicians, containing demographic, socio-economic, health assistance and cause of death data. Automated selection of underlying cause of death is in use in Brazil since 1996 at state and municipal health departments, following International Classification of Diseases 10^th^ Revision (ICD10) coding rules for selection and modification [7]. Both underlying and associated causes registered in parts I and II of the international form of medical certificate of cause of death are recorded in SIM since 1996. Associated causes included both consequential (terminal and intervening) and contributing causes of death [8]. All causes registered in death certificates, including symptoms, signs and modes of dying were included.

Non foetal death records of residents in the state of Amazonas that occurred within the State from January 1^st^ 2006 until December 31^st^ 2014 were included in the study after eliminating all personal identifiers. Multiple cause of death data was analysed under a person-based approach, eliminating duplication of counts of reported causes for each death [4]. Death records were classified based on tuberculosis reporting in any part of the death certificate corresponding to ICD10 codes A15-A19 (Tuberculosis block of three-character categories), B90 (Sequelae of tuberculosis three-character category) and B20.0 (HIV disease resulting in mycobacterial infection four-character subcategory) without mention to the former, as follows: tuberculosis not reported (TBNoR), tuberculosis reported as the underlying cause of death (TBUC) and tuberculosis reported as an associated cause of death (TBAC).

Age standardized annual mortality rates for TBUC, TBAC and with TB reported (TBUC plus TBAC) were estimated for the State of Amazonas, using the direct standardization method and WHO 2000-2025 standard population [9].

Associated and underlying causes related, respectively, to TBUC and TBAC deaths were listed in tabulation lists according to tuberculosis natural history of disease and frequency criteria.

Mortality odds ratios (OR) of reporting TBUC and TBAC in relation to TBNoR were estimated using multinomial logistic regression modelling [10]. Covariates were added in order to adjust for potential confounding as follows: triennium of death (2006-08, 2009-11 and 2012-14), place of residence prior to, and of occurrence of death (state capital – Manaus – and all other municipalities), sex (male, female and ignored), age group (0-14, 15-39, 40-64, 65 and more years and ignored), race/skin color (white, nonwhite and ignored) and education level (0-7, 8-11, 12 and more school years, not applicable and ignored). Nonwhite race/skin color comprised all categories available in the death certificates except white, including black, yellow, mulatto and indigenous. Education level was classified as not applicable for all children under six years of age according to Brazilian education policy. Records without information on covariates (ignored), interpreted as proxies of lack of data quality, were kept in the models in order to investigate association with TB reporting.

The categories of the covariates were inserted as indicator (dummy) variables [11]. Statistical significance was set at the 20% and 5% levels, respectively, for the entry and retention of covariates in the model, and 95% confidence limits were calculated for final model adjusted OR estimates. Statistical significance was established by the Wald test and the quality of fit of the final model by the analysis of deviance measures [10]. Analyses were developed using Stata 12 (StataCorp. 2011. *Stata Statistical Software: Release 12*. College Station, TX: StataCorp LP).

In order to investigate potential bias related to single cause reporting due to ill-defined underlying causes, alternative models were fitted excluding deaths due to ICD10 codes R98 (Unattended death) and R99 (Other ill-defined and unspecified causes of mortality) in the TBNoR group.

The study was approved by the Ethics in Research Board Committee of the Alfredo da Mata Hospital Foundation, Amazonas, under the National Ethics in Research Board Committee - CONEP (protocol number – CAAE - 59417016.3.0000.0002; available at http://plataformabrasil.saude.gov.br/login.jsf).

## Results

Tuberculosis was reported in 1,933 (1.6%) out of 120,701 death certificates registered in the state of Amazonas over the years 2006 to 2014. Tuberculosis was assigned as the underlying cause in 1,167 (60.4%) death certificates, 1.5 times more frequently than as an associated cause (766; 39.6%). All forms of active tuberculosis (A15-A19), the most frequent way of reporting (1.20%; n = 1,445), were 2.7 times more likely to be coded as underlying cause of death. Sequelae of tuberculosis (B90) was reported in 189 (0.16%) death certificates, 1.6 times more likely as the underlying cause. HIV disease resulting in mycobacterial infection (B20.0), without mention to ICD10 codes A15-A19 or B90, was found in 309 death records (0.26%), always as associated cause. Among deaths with tuberculosis reported as the underlying cause of death (1,167), 949 (81.3%) were due to tuberculosis of the respiratory tract, 31 (2.6%) of the nervous system, 28 (2.4%) of all other organs, 45 (3.9%) to miliary tuberculosis and 114 (9.8%) to sequelae of tuberculosis.

Age standardized annual TBUC, TBAC and TB reported mortality rates are presented in figure 1. TBUC mortality ranged from 5.9 to 7.8/100,000, peaking in 2009 and remaining below 7.0/100,000 thereafter. TBAC mortality varied from 2.7 to 4.0/100,000 – approximately half of TBUC mortality range – peaking in 2010 (4.0/100,000) and 2013 (3.9/100,000). Overall mortality with TB reported ranged between 8.9 and 11.1/100,000, exceeding 10/100,000 only in 2009 (11.1/100,000) and 2013 (10.8/100,000).

**Figure 1.**
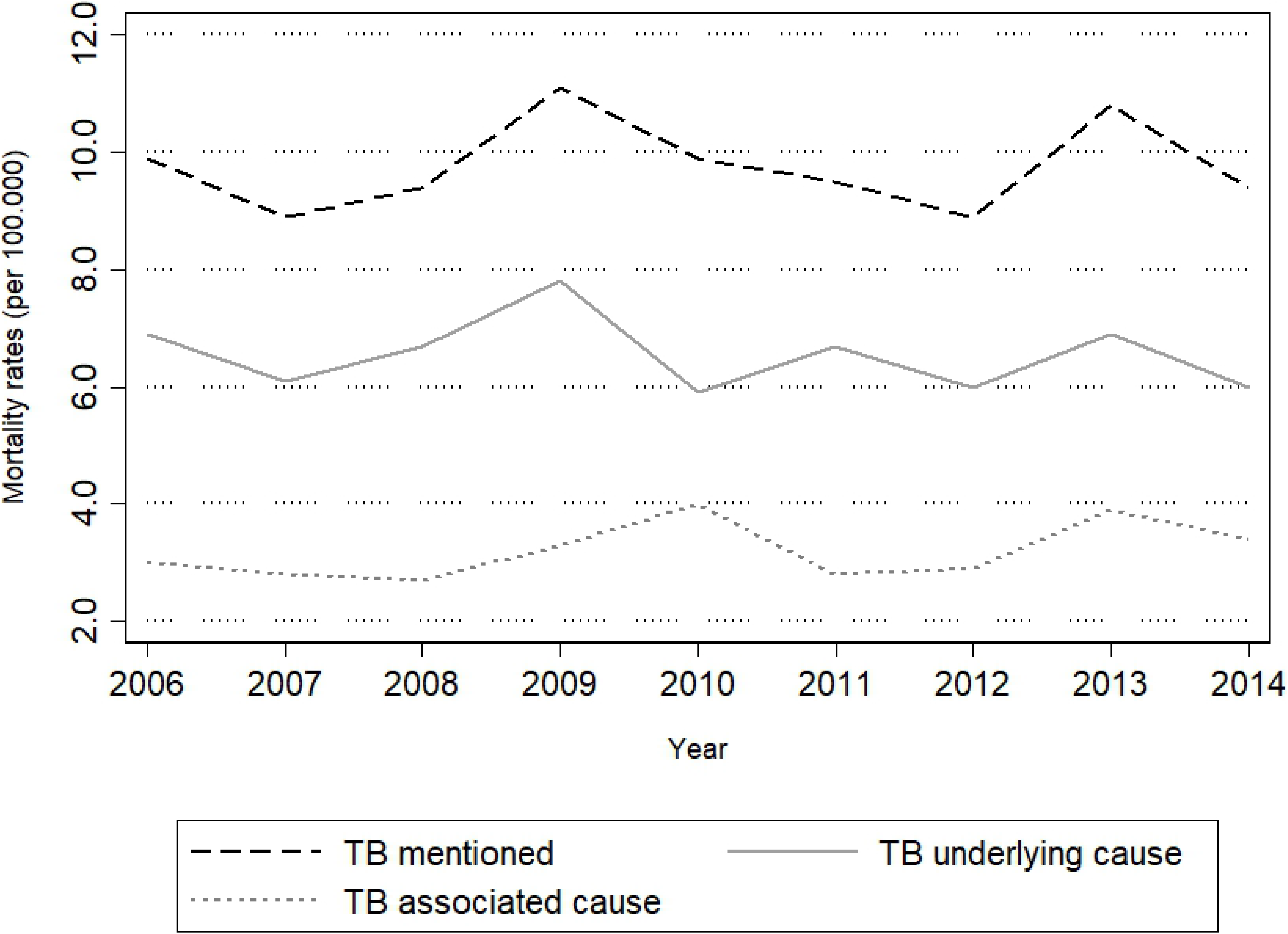
Age adjusted annual mortality rates of tuberculosis as underlying cause, associated cause and with tuberculosis mentioned in death certificates from 2006 to 2014, State o Amazonas, Brazil

Most of the associated causes mentioned with tuberculosis reported as the underlying cause (Table 1 – left side) may be interpreted as terminal, such as respiratory failure and sepsis – mentioned in 30% or more of death certificates – and renal failure. Other causes include pneumonias, nutritional and metabolic disorders – predominantly malnutrition and diabetes mellitus -, diseases of the respiratory, cardiovascular and digestive systems – chronic lower respiratory, lung insterstitium, hypertensive and liver diseases - and mental disorders - psychoactive substance use, mainly alcohol and tobacco use (data not shown).

**Table 1.**
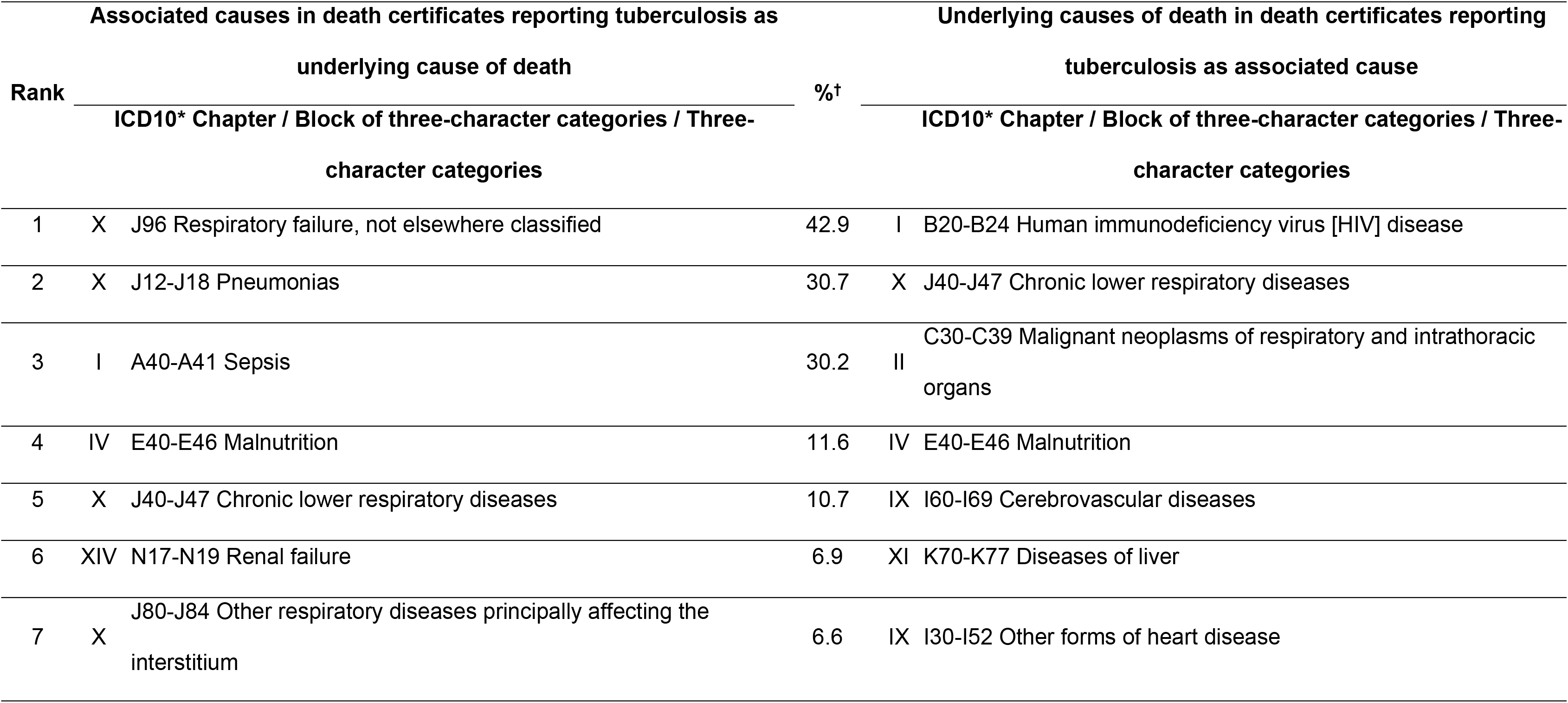
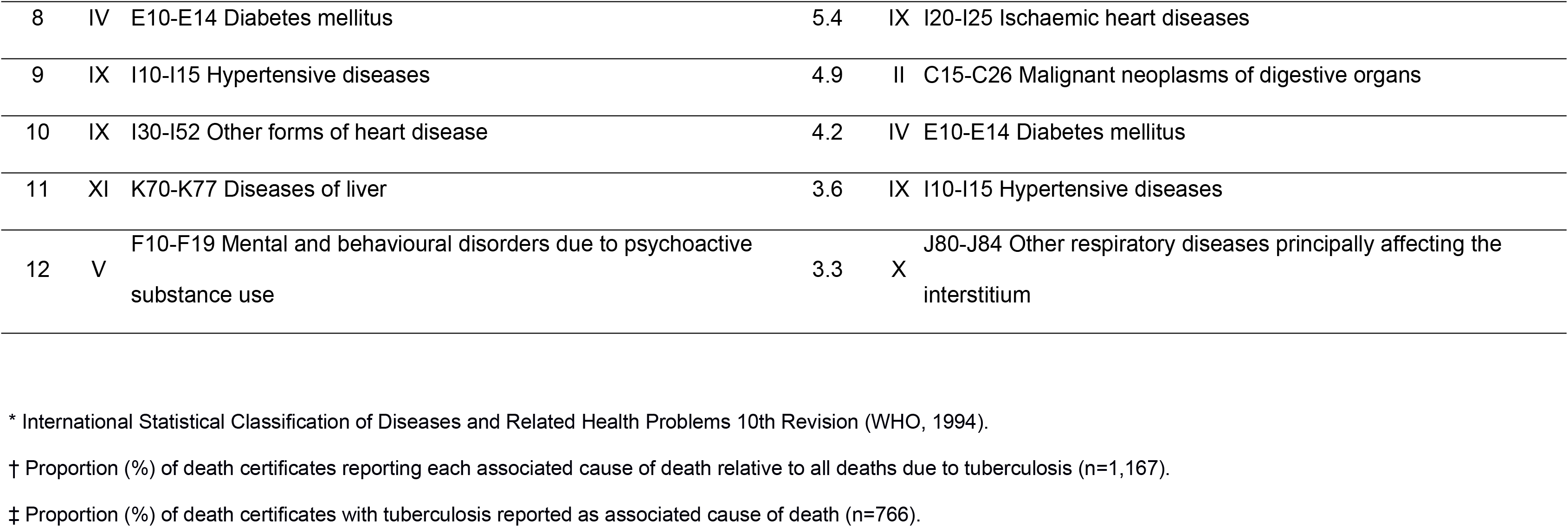
Distribution (%) of associated causes in death certificates reporting tuberculosis as underlying cause of death, and underlying causes in death certificates reporting tuberculosis as associate cause of death, Amazonas State, Brazil, 2006 to 2014.

| Associated causes in death certificates reporting tuberculosis as |  |  |  | Underlying causes of death in death certificates reporting |  |  |  |
| --- | --- | --- | --- | --- | --- | --- | --- |
| Rank | underlying cause of death |  |  | %† | tuberculosis as associated cause |  |  |
|  | ICD10* Chapter / Block of three-character categories / Three-character categories |  |  |  | ICD10* Chapter / Block of three-character categories / Three-character categories |  |  |
| 1 | X | J96 | Respiratory failure, not elsewhere classified | 42.9 | I | B20-B24 | Human immunodeficiency virus [HIV] disease |
| 2 | X | J12-J18 | Pneumonias | 30.7 | X | J40-J47 | Chronic lower respiratory diseases |
| 3 | I | A40-A41 | Sepsis | 30.2 | II | C30-C39 | Malignant neoplasms of respiratory and intrathoracic organs |
| 4 | IV | E40-E46 | Malnutrition | 11.6 | IV | E40-E46 | Malnutrition |
| 5 | X | J40-J47 | Chronic lower respiratory diseases | 10.7 | IX | I60-I69 | Cerebrovascular diseases |
| 6 | XIV | N17-N19 | Renal failure | 6.9 | XI | K70-K77 | Diseases of liver |
| 7 | X | J80-J84 | Other respiratory diseases principally affecting the interstitium | 6.6 | IX | I30-I52 | Other forms of heart disease |

|  |  |  |  |  |  |
| --- | --- | --- | --- | --- | --- |
| 8 | IV | E10-E14 Diabetes mellitus | 5.4 | IX | I20-I25 Ischaemic heart diseases |
| 9 | IX | I10-I15 Hypertensive diseases | 4.9 | II | C15-C26 Malignant neoplasms of digestive organs |
| 10 | IX | I30-I52 Other forms of heart disease | 4.2 | IV | E10-E14 Diabetes mellitus |
| 11 | XI | K70-K77 Diseases of liver | 3.6 | IX | I10-I15 Hypertensive diseases |
| 12 | V | F10-F19 Mental and behavioural disorders due to psychoactive substance use | 3.3 | X | J80-J84 Other respiratory diseases principally affecting the interstitium |

\* International Statistical Classification of Diseases and Related Health Problems 10th Revision (WHO, 1994).
† Proportion (%) of death certificates reporting each associated cause of death relative to all deaths due to tuberculosis (n=1,167).
‡ Proportion (%) of death certificates with tuberculosis reported as associated cause of death (n=766).

When tuberculosis was an associated cause, seven of each ten deaths were due to infectious diseases, almost exclusively to HIV disease (Table 1 – right side). Respiratory diseases, neoplasms and cardiovascular diseases together accounted for approximately 21% of deaths, mainly due to, respectively, chronic lower respiratory diseases (6.3%), cancer of respiratory and intrathoracic organs (2.5%) and hypertension related diseases (cerebrovascular, ischemic and hypertensive diseases – 4.3%). Other relevant causes include malnutrition, liver diseases and diabetes mellitus, together accounting for 5% of deaths.

Seven of the top twelve causes listed in either side Table 1 – namely malnutrition, diabetes mellitus, chronic lower and insterstitium related respiratory diseases, hypertensive and other forms of heart diseases, and liver diseases - were assigned in death certificates reporting tuberculosis indistinctively as underlying or associate cause of death.

Adjusted OR of reporting TBUC (against TBNoR) were significantly lower among deaths of residents in the state capital (34% decrease), women (13% decrease) and with increasing education level (33% and 50% decrease, respectively, with 8 to 11 and 12 or more school years). When education level did not apply (among children aged 0 to 5 years) the chance of tuberculosis as the underlying cause decreased 86%. Higher odds of TBUC was observed when deaths occurred among non-whites (38% increase) and in the state capital (69% increase) (Table 2).

**Table 2.**
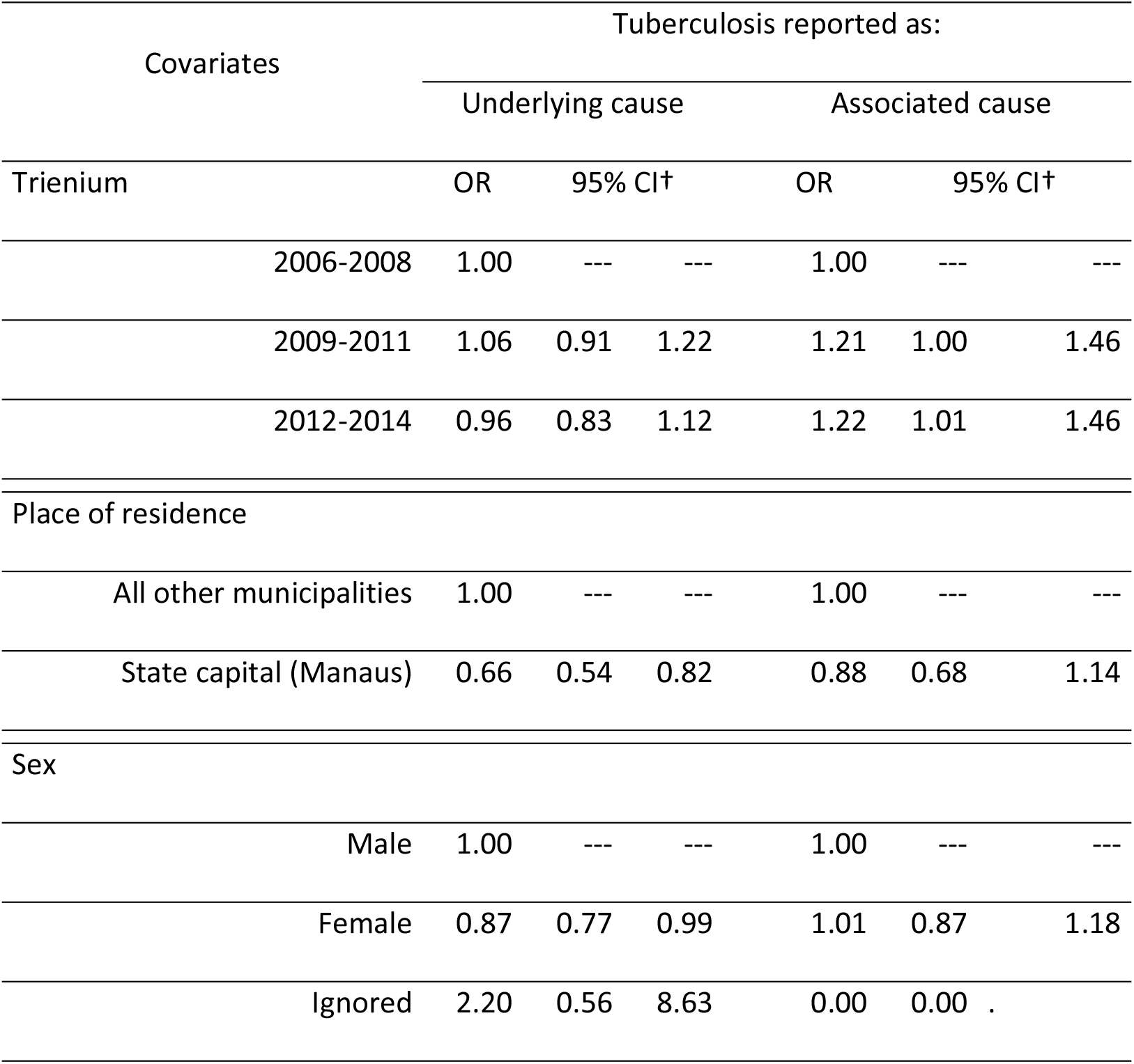
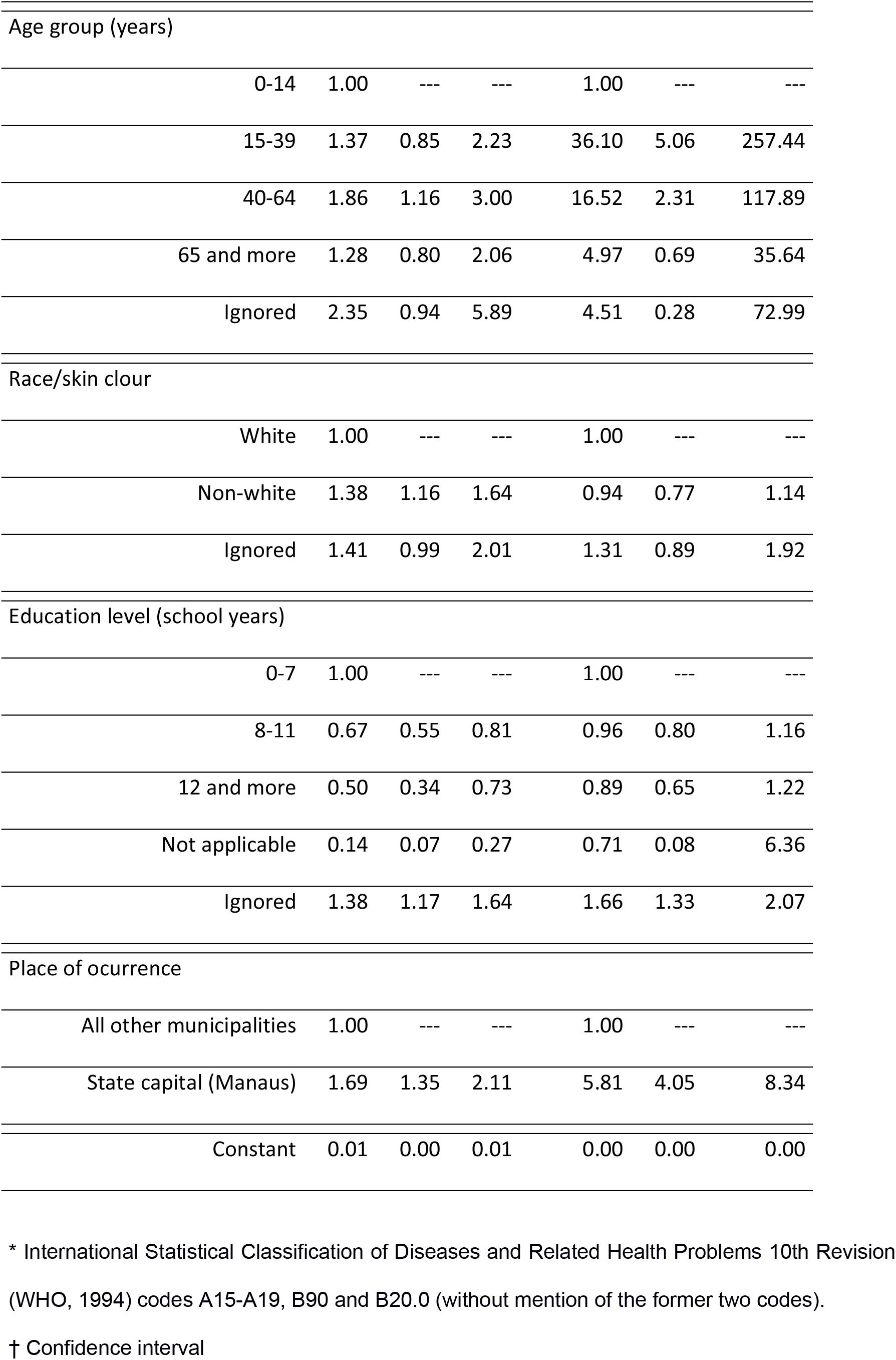
Adjusted odds ratios of reporting tuberculosis* as underlying and associate cause of death, State of Amazonas, Brazil, 2006 to 2014

The adjusted OR of reporting tuberculosis as associated cause increased in the last two trienniums (21% and 22%, respectively, in years 2009 to 2011 and 2012 to 2104), with adult ages (36.1 and 16.5 times higher, respectively, within ages 15 to 39 and 40 to 64 years) and when death occurred in the state capital (5.8 times higher).

No significant changes were noted following alternative models excluding deaths due to single cause ICD10 three-character categories R98 (Unattended death) and R99 (Other ill-defined and unspecified causes of mortality) in the TBNoR group (data not shown).

## Discussion

Although predominantly assigned as the underlying cause, tuberculosis reporting as associate cause corresponded to 40% of all mentions of tuberculosis mortality in the State of Amazonas from 2006 to 2014. Deaths with tuberculosis as associated cause are not usually counted in primary statistics of mortality. Considering all death certificated in which tuberculosis was mentioned, an average 50% increase in annual tuberculosis related mortality rates was observed when compared with TBUC mortality rates. Similar patterns of tuberculosis reporting in death certificates were observed elsewhere in Brazil [5,12–14].

Causes associated with tuberculosis were mainly respiratory, infectious, nutritional, metabolic, cardiovascular and digestive diseases – frequently terminal diseases. Such causes are among those characterized as complications of tuberculosis. Improvements in the diagnosis and treatment of tuberculosis patients in a timely manner are among the main measures necessary to avoid the occurrence of these outcomes [15].

The high frequency of tuberculosis related mortality associated with malnutrition, diabetes mellitus, alcoholism and smoking found in the present study are in accordance with reports in the scientific literature [16]. These findings reflect the role of emerging risk factors such as chronic diseases, socio-economic and behavioral aspects in increasing the unsuccessful TB treatment outcomes. Knowledge of these causes is important in guiding control and preventing tuberculosis-related death [16].

Mortality due to tuberculosis was associated with poor socioeconomic conditions (living outside Manaus, non-white race/skin color and low education level) and negatively related to female sex. The higher odds of dying due to tuberculosis among non-whites, including natives of indigenous origin, and its inverse relation with education level may reflect health related social inequalities. These findings corroborate literature data that showed the relationship between TB and socioeconomic deprivation and ethnic status in the state of Amazonas [17]. Similarly, the lower odds of tuberculosis reporting as the underlying cause observed among deaths of residents in the State capital may be explained by higher socioeconomic level and health care access of the population living in Manaus compared to the remaining municipalities. The association of male sex and poor socioeconomic conditions with high tuberculosis mortality has also been observed in other populations [18–20].

Mortality with tuberculosis was positively associated with adult ages, reflecting its strong relation as associate cause with HIV disease [14]. HIV disease was by far the most frequent underlying cause of deaths in which tuberculosis was reported as an associate cause, followed by diseases of the respiratory and cardiovascular systems, neoplasms, nutritional and metabolic disorders and digestive diseases. Similar cause specific mortality profiles were found in two cohort studies involving tuberculosis patients in Brazil and China [13,21].

The high frequency of deaths due to AIDS with mention of tuberculosis may reflect the natural history of HIV disease in high tuberculosis prevalence settings, as in the State of Amazonas. However, it may also suggest inadequate implementation of HIV disease control Brazilian guidelines, such as early HIV infection diagnosis, latent tuberculous infection screening and 6-month isoniazid preventive therapy among HIV patients. In Brazil HIV disease is commonly diagnosed following tuberculosis onset and the association of lack of isoniazid preventive therapy and tuberculosis incidence has been observed among HIV patients [22].

The increasing odds of tuberculosis reporting as associate cause with recent years suggests an overall improvement in diagnosis, more specifically of HIV disease and related comorbidities. In a recent study the linkage of tuberculosis and HIV/Aids case report databases reduced underreporting and improved the epidemiological surveillance of the two diseases in the state of Amazonas [23]. Also, quality of cause of death assignment has improved in Amazonas, with decreasing trends in mortality due to ill-defined and unspecified causes [24].

The higher odds of reporting tuberculosis when death occurred at the State capital, even higher as associate cause, may reflect the concentration of health care facilities in Manaus, including reference health services for diagnosis and treatment of infectious and non-infectious diseases. This hypothesis is supported by studies having shown that the state capital presents better levels of performance of the health services [23,25].

Tuberculosis remains a significant burden in developing countries. Notwithstanding all efforts aimed at its control and the increasing coverage of primary health care units in Brazil from 2006 to 2014, high incidence rates are expected for future decades as a consequence of high prevalence of *Mycobacterium tuberculosis* infection following present and past high transmission rates. However, deaths due to tuberculosis are, in most, avoidable once access to prompt and adequate health care is available [13].

Among the limitations of our study, cause of death records, the only data source used, may lack sensitivity and specificity required for medical outcomes, more specifically for tuberculosis diagnosis [13,26]. Although systematically improving along the past decades, SIM still lacks completeness and quality of cause of death information in Brazil, especially in the Northern regions - including the State of Amazonas – where proportional mortality due to ill-defined causes, although presenting a declining trend, remain high [27].

Despite such limitations, our findings are consistent with other authors based on studies developed in different settings, time periods and with distinct populations. Also, predictors of tuberculosis reporting in death certificates remained unchanged after excluding deaths due to ICD10 codes for ill-defined causes.

## Conclusions

Tuberculosis mortality was predominantly reported as the underlying cause of death in the State of Amazonas, from 2006 to 2014. However, a substantial increase (50%) in mortality rate estimates followed the inclusion of death certificates in which tuberculosis was assigned as associate cause.

Tuberculosis was concurrently reported in death certificates with a myriad of infectious and non-infectious diseases, as well as with ill-defined causes. Causes associated with tuberculosis assigned as underlying cause of death were mainly related to clinical complications and exposure to risk factors. HIV disease was the main underlying cause of deaths in which tuberculosis appeared as associate cause.

Tuberculosis reporting as the underlying cause of death was associated with male gender and indicators of unfavourable socioeconomic conditions and health care access constraints, whereas reporting as associate cause was related to adult ages, typical of high HIV disease incidence, and occurrence in recent years, due to improvements in health care and cause of death assignment. The occurrence of death in Manaus, the State capital, positively predicted both types of tuberculosis mortality reporting – more strongly as associated cause – possibly due higher coverage and better performance of health care and diagnostic facilities.

## Author contributions

Wrote the paper: VSS AJLC DBC. Data analysis and interpretation: VSS LCFS PCSB AJLC. Data acquisition: PCSB AAC. Critical revision of the manuscript for important intellectual content: VSS LCFS DBC AAC AJLC. Manuscript concept and design: VSS AJLC.

